# Population Turnover in Remote Oceania Shortly After Initial Settlement

**DOI:** 10.1101/268037

**Authors:** Mark Lipson, Pontus Skoglund, Matthew Spriggs, Frederique Valentin, Stuart Bedford, Richard Shing, Hallie Buckley, Iarawai Phillip, Graeme K. Ward, Swapan Mallick, Nadin Rohland, Nasreen Broomandkhoshbacht, Olivia Cheronet, Matthew Ferry, Thomas K. Harper, Megan Michel, Jonas Oppenheimer, Kendra Sirak, Kristin Stewardson, Kathryn Auckland, Adrian V.S. Hill, Kathryn Maitland, Stephen J. Oppenheimer, Tom Parks, Kathryn Robson, Thomas N. Williams, Douglas J. Kennett, Alexander J. Mentzer, Ron Pinhasi, David Reich

## Abstract

Ancient DNA analysis of three individuals dated to ~3000 years before present (BP) from Vanuatu and one ~2600 BP individual from Tonga has revealed that the first inhabitants of Remote Oceania (“First Remote Oceanians”) were almost entirely of East Asian ancestry, and thus their ancestors passed New Guinea, the Bismarck Archipelago, and the Solomon Islands with minimal admixture with the Papuan groups they encountered [1]. However, all present-day populations in Near and Remote Oceania harbor 25-100% Papuan ancestry, implying that there must have been at least one later stream of migration eastward from Near Oceania. We generated genome-wide data for 14 ancient individuals from Efate and Epi Islands in Vanuatu ranging from 3,000-150 BP, along with 185 present-day Vanuatu individuals from 18 islands. We show that people of almost entirely Papuan ancestry had arrived in Vanuatu by 2400 BP, an event that coincided with the end of the Lapita cultural period, changes in skeletal morphology, and the cessation of long-distance trade between Near and Remote Oceania [2]. First Remote Oceanian ancestry subsequently increased via admixture but remains at 10-20% in most islands. Through a fine-grained comparison of ancestry profiles in Vanuatu and Polynesia with diverse groups in Near Oceania, we find that Papuan ancestry in Vanuatu is consistent with deriving from the Bismarck Archipelago instead of the geographically closer Solomon Islands. Papuan ancestry in Polynesia also shows connections to the ancestry profiles present in the Bismarck Archipelago but is more similar to Tolai from New Britain and Tutuba from Vanuatu than to the ancient Vanuatu individuals and the great majority of present-day Vanuatu populations. This suggests a third eastward stream of migration from Near to Remote Oceania bringing a different type of Papuan ancestry.

## Results and Discussion

We generated genome-wide data for 14 ancient individuals from Central Vanuatu (**Table 1**; **Table S1**). Of these, 11 individuals are newly reported, and 3 individuals that were previously published are represented here by higher quality data [1]. We identified and selected cochlear bone sections of petrous bones and processed them into powder in dedicated clean rooms at University College Dublin [3]. We then shipped the powder to Harvard Medical School, where in a second set of clean rooms we extracted DNA [4, 5] and created individually barcoded Illumina sequencing libraries, some of which we treated with the enzyme Uracil-DNA Glycosylase (UDG) to greatly reduce the characteristic errors associated with degraded ancient DNA [6, 7]. We screened these libraries for evidence of authentic ancient DNA by enriching for DNA overlapping the mitochondrial genome [8], sequencing on an Illumina NextSeq500 instrument, and assessing the data based on rates of cytosine-to-thymine damage in the terminal nucleotide and consistency with the consensus mitochondrial genome (STAR Methods) [9]. For libraries that were promising after screening, we enriched for regions targeting approximately 1.24 million single nucleotide polymorphisms (SNPs) in the human genome and sequenced the enriched products to greater depth (STAR Methods). We determined sex by examining the ratio of sequences overlapping the Y chromosome and X chromosome, and for males, we additionally estimated contamination based on the rate of polymorphism on the haploid X chromosome (STAR Methods; **Table S1**). The data for the 14 individuals passing quality control were derived from a total of 46 Illumina libraries (1-8 per individual; **Table S2**). We also generated genome-wide SNP genotype data on the Human Origins array for 185 present-day individuals from Vanuatu who gave informed consent for studies of genetic variation, with approval from both the University of Oxford and the Vanuatu Cultural Centre (STAR Methods; **Table S3**).

**Table 1.** Details of Ancient Vanuatu Samples Analyzed in this Study

| Sample | Code | Date | Population label | Location | Country | Sex | mtDNA | Y | SNPs |
| --- | --- | --- | --- | --- | --- | --- | --- | --- | --- |
| I1370 | B17.P3 | 1160-830 calBCE (3083±26 BP, Wk-21026, corrected for Marine Reservoir Effect) | Vanuatu_3000BP | Teouma, Efate | Vanuatu | F | <u>B4a1a1</u> |  | 237405 |
| I1369 | B10B.P3 | 1050-800 calBCE (3045±30 BP, Poz-81126, corrected for Marine Reservoir Effect) | Vanuatu_3000BP | Teouma, Efate | Vanuatu | F | <u>B4a1a</u> |  | 271048 |
| I1368 | B30A.P3 | 1040-790 calBCE (2983±32 BP, Wk-22657, corrected for Marine Reservoir Effect) | Vanuatu_3000BP | Teouma, Efate | Vanuatu | F | <u>B4a1a</u> |  | 185282 |
| I5951 | TeoQE | 1258-1088 calBCE (2955±20, PSUAMS-2411) | Vanuatu_3000BP | Teouma Quarry Edge | Vanuatu | M | <u>B4a1a1</u> |  | 23107 |
| I4451 | TAP1 | 516-369 calBCE (2348±32 BP, Wk-20390) | Vanuatu_2400BP | Mele-Taplins, Efate | Vanuatu | M | M28a | K2b1 | 340152 |
| I4425 | EF3_2_E | 1652-1950 calCE (200±20 BP, UCIAMS-188795) | Vanuatu_150BP | Ifira, Efate | Vanuatu | F | P2 |  | 700783 |
| I4450 | SEPU1 | 1520-1645 calCE (305±15 BP, UCIAMS-188793) | Vanuatu_350BP | Pangpang, Efate | Vanuatu | F | P1d2 |  | 735460 |
| I4096 | BURU5B | 429-595 calCE (1430±20 BP, PSUAMS-1841) | Vanuatu_1400BP | Burumba, Epi Island | Vanuatu | M | <u>B4a1a1k</u> | K2b1 | 888003 |
| I3921 | BURU5D | 400-700 CE [429-595 calCE (1430±20 BP, PSUAMS-1841) from burial 5 skull B; 551-650 CE (1464±30 BP, Wk-25769)] | Vanuatu_1400BP | Burumba, Epi Island | Vanuatu | M | P1d1 | K2b1 | 855305 |
| I5259 | Mang1 | 1307-1430 calCE (559±30 BP, Wk-20030) | Vanuatu_600BP | Mangaliliu | Vanuatu | F | P1f |  | 799098 |
| I4419 | BB1 | 1678-1940 calCE (135±15 BP, UCIAMS-188792) | Vanuatu_150BP | Banana Bay, Efate | Vanuatu | M | <u>B4a1</u> | K2b1 | 763556 |
| I4424 | EF_Pango1 | 1661-1950 calCE (190±15 BP, UCIAMS-188794) | Vanuatu_150BP | Pango Village, Efate | Vanuatu | M | *R* | M1b | 780469 |
| I4105 | WAMB1 | 1529-1798 calCE (255±20 BP, PSUAMS-1922) | Vanuatu_200BP | Wambi Bay, Epi Island | Vanuatu | M | M28a | <u>O1a2</u> | 1012081 |
| I4106 | WAMB2 | 1645-1950 calCE (225±20 BP, PSUAMS-1923) | Vanuatu_200BP | Wambi Bay, Epi Island | Vanuatu | M | <u>B4a1a1a11</u> | <u>O1a2</u> | 1020436 |

Note: Underlining indicates typical East Asian (First Remote Oceanian) haplogroups, while lack of underlining indicates typical Australo-Papuan haplogroups (the italicized mtDNA haplogroup *R* is unclassified). The first three samples listed are previously published individuals [1] but with new libraries now added to increase coverage; the other 11 are newly published individuals.

## Clustering analyses

We performed automated clustering analysis with the ADMIXTURE software [10], using a data set consisting of the ancient and present-day Vanuatu samples together with other Oceanian, East Asian, and worldwide populations genotyped on the Human Origins array [1] (**Figure 1**; **Figure S1**). At K = 8 clusters, four ancestry components were inferred to be widespread in Oceania. Three correlate (predominantly) to Papuan ancestry, and are maximized in New Guinea (purple in the ADMIXTURE plot), Mamusi and Baining from New Britain (blue), and Nasioi from Bougainville in the Solomon Islands (red). The fourth component (green) correlates to First Remote Oceanian ancestry, and is maximized in the ancient Lapita individuals from Vanuatu and Tonga. Other Oceanian populations display variable combinations of these components, forming gradients of ancestry between New Guinea, New Britain and New Ireland in the Bismarck Archipelago, and the Solomon Islands. The great majority of present-day as well as ancient groups from Vanuatu show highly similar ratios of the three Papuan ancestry components (although their First Remote Oceanian proportions vary), suggesting that they largely derived their Papuan ancestry from the same source. Among populations in Near Oceania, the most similar to Vanuatu in terms of the Papuan ancestry component ratio (purple-to-blue-to-red) are groups from New Britain in the Bismarck Archipelago with a majority of the blue component and smaller contributions of purple and red, pointing to an origin from the Bismarck Archipelago (rather than the geographically closer Solomon Islands) for the Papuan ancestry in Vanuatu. A similar pattern was previously inferred for the origin of the Papuan ancestry in Santa Cruz to the north of Vanuatu [11] (a result we replicate here), implying similar sources for both island chains.

**Figure 1.**
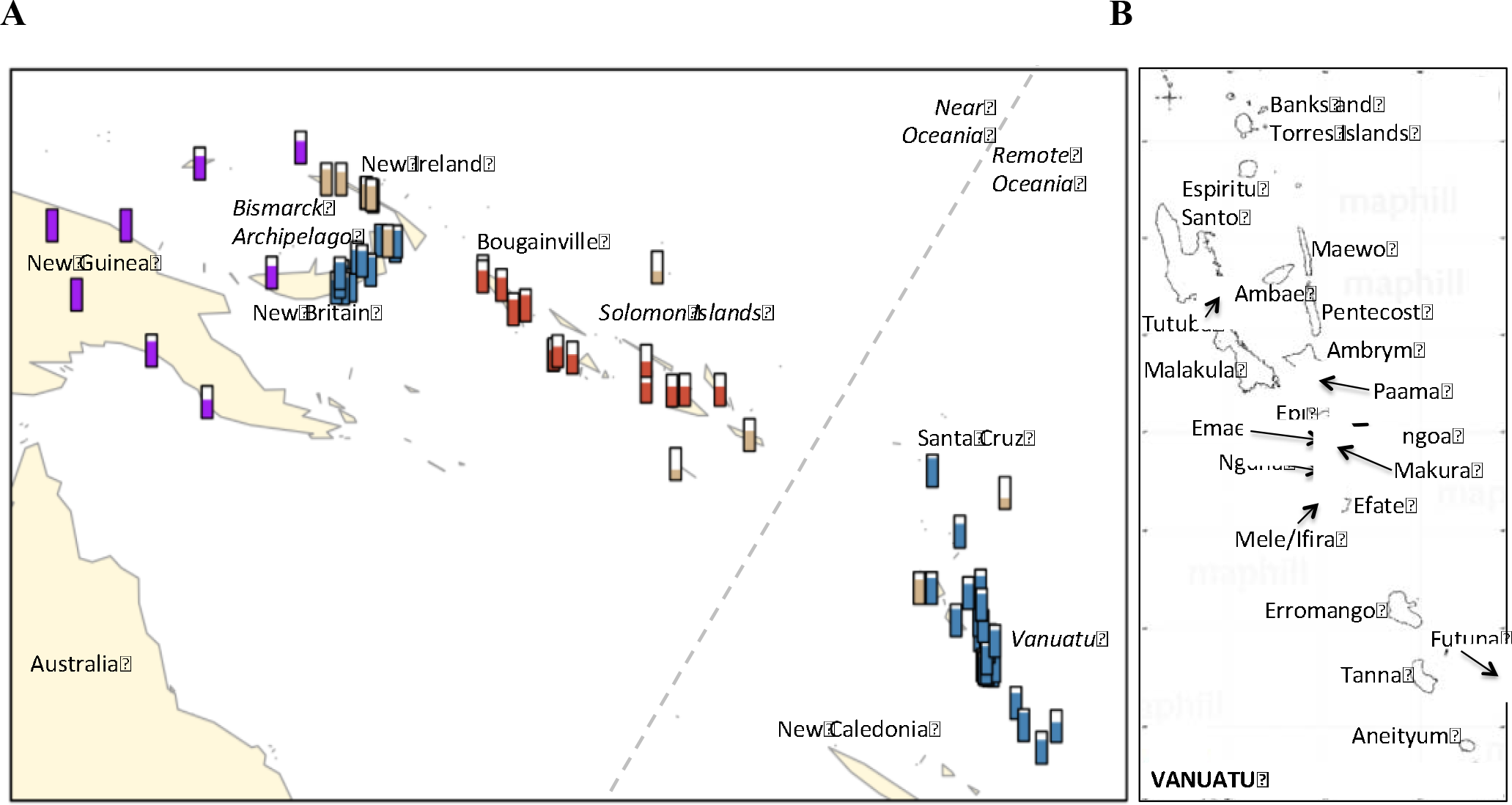
Locations and broad-scale genetic structure of analyzed populations. (A) Bars represent proportions of Papuan and First Remote Oceanian (white) ancestry. Purple, red, and blue and colors match those in **Figure S1** but here correspond to clusters assigned based on the proximity of populations in the ADMIXTURE results (i.e., overall ratios of Papuan ancestry components) rather than individual ADMIXTURE components: purple, similar to the ratio maximized in New Guinea; blue, similar to New Britain; red, similar to Solomon Islands; brown, mixed between New Britain and Solomon Islands clusters (primarily New Ireland). (B) Map of Vanuatu with islands labeled from which ancient or present-day data are reported in this study. Map data are from freely available sources: (A) was plotted in R using the ‘maps’ package with data from http://www.naturalearthdata.com/, and (B) was made with a blank map downloaded from http://www.maphill.com/vanuatu/simple-maps/blank-map/no-labels/.

We also carried out a principal component analysis focusing on the geographic variation in Papuan ancestry (**Figure S2**). The results confirm those from ADMIXTURE, with the primary feature being a U-shaped cline from top left to top right—encompassing Nakanai (western New Britain), Sulka and Mengen (eastern New Britain), most of Vanuatu, Tolai, Tutuba, New Ireland, and finally Bougainville—corresponding closely to a trend of increasing red and decreasing blue components in ADMIXTURE. The position of the Vanuatu samples in the PCA also supports the hypothesis that the inhabitants of the region after the initial Lapita settlement derived ancestry ultimately not from the closer Solomon Islands but from the area of New Britain in the Bismarck Archipelago.

## Papuan and First Remote Oceanian ancestry proportions

It has been shown that the strongest driver of genetic variation in Oceania today is the widespread but highly variable admixture between Papuan and First Remote Oceanian ancestry sources, the former representing original inhabitants of Near Oceania and the latter descendants of the Austronesian expansion from East and Southeast Asia [1]. From our clustering results, a dramatic turnover is apparent in Vanuatu between around 3000 and 2400 years ago, with First Remote Oceanian populations being joined or possibly completely replaced by individuals of (almost) entirely Papuan ancestry. To provide precise estimates of mixture proportions, we used *f*_*4*_-ratio statistics [12], assuming a topology of (Atayal, (Kankanaey, First Remote Oceanian)) for East Asian-derived ancestry and (Australian, (New Guinea, Papuan)) for Papuan ancestry (**Figure 1**; **Table S4**). Taking advantage of our increased coverage compared to the first study of Lapita samples, we find that the ~3000 BP Lapita individuals likely had a small amount of Papuan-related ancestry (2.4 ± 0.9%), although it remains striking that the initial First Remote Oceanian migrants were only minimally admixed. Given the small proportion, we did not have sufficient statistical power to determine whether this Papuan-related ancestry is derived from the region surrounding New Guinea or could perhaps have been acquired elsewhere, such as in the Philippines or eastern Indonesia. Notably, the first post-Lapita sample (2400 BP from Mele-Taplins, Efate) had almost entirely Papuan ancestry but with a small amount from First Remote Oceanians (4.2 ± 1.1%). The more recent ancient individuals are similar in their proportions to present-day populations: 8-12% First Remote Oceanian ancestry for 1400-200 BP and 20% for 150 BP (Efate), as compared to a range of 9-38% today (mostly 12-20%; maximized in the Polynesian outlier population of Futuna). For time points with multiple samples, the individuals’ mixture proportions are statistically indistinguishable, except at 150 BP (~14%, 21%, and 26% First Remote Oceanian).

## Dates of admixture

We estimated dates of admixture based on weighted admixture linkage disequilibrium (LD) [13] using ALDER [14], with Ami and New Guinea as references (**Figure 2**; **Table S4**). We obtain significant evidence for admixture LD in almost all present-day populations and three ancient population groupings (noting that power is highly sample size-dependent). The date estimates are mostly 40-100 generations ago, or 1,100-2,800 years ago assuming 28 years per generation [15], consistent with initial admixture soon after the early settlement of Vanuatu and further mixture continuing through time (in cases of multiple pulse of admixture, ALDER produces a single average date). We observe a modest but significant negative correlation between admixture date and First Remote Oceanian ancestry proportion (R^2^ = 0.32 for populations in **Figure 2**, nominal p < 0.01), as would be expected if a subset of populations (e.g., Efate, Emae, Futuna, Makura) received more recent pulses of gene flow from groups with high proportions of First Remote Oceanian ancestry (a plausible scenario in light of Polynesian cultural influence [16]). We also obtain a direct admixture date of 18 ± 6 generations in the past (500 ± 160 years) for a pair of ancient samples from Vanuatu radiocarbon dated to ~1,400 years ago, consistent with the ALDER dates in the majority of present-day groups. There has been debate about the timing of admixture between people of East Asian and Papuan ancestry in Remote Oceania, with methods based on wavelet transformations suggesting mixing >3,000 BP, prior to the Lapita expansion to Remote Oceania [11, 17], and methods based on admixture LD suggesting more recent dates, implying that mixture must have occurred following later streams of gene flow [18]. It was recently argued that the differences may reflect systematic biases of the methods for dates more than a couple of thousand years old [11], and thus our finding of a definitively post-Lapita date in samples that are within a thousand years of the estimated admixture date strengthens the evidence for more recent mixture.

**Figure 2.**
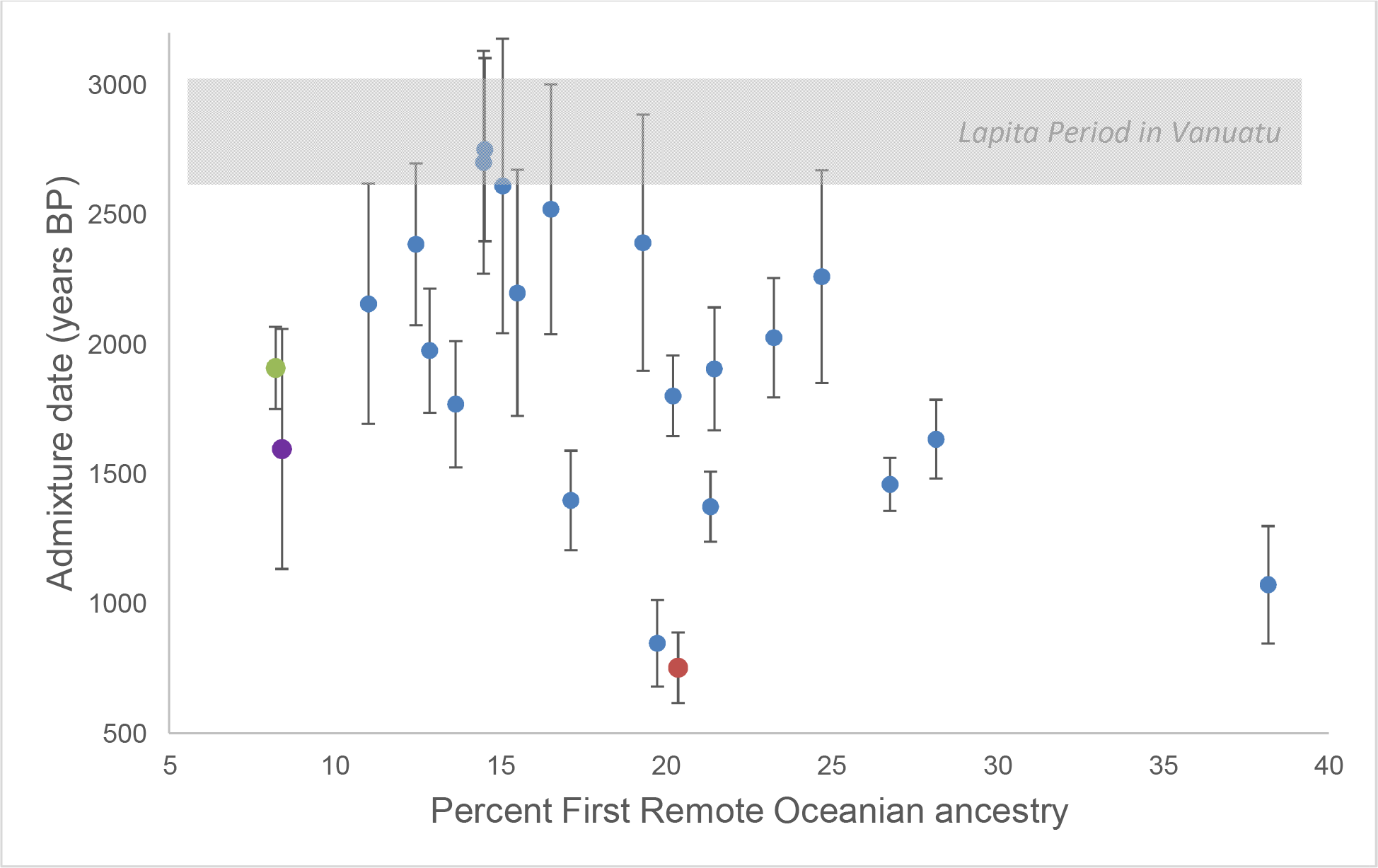
Ancestry proportions and dates of admixture in Vanuatu. Blue points represent the 0 present-day populations with the most confident admixture date estimates (as measured by Z-score for difference from zero). Colored points represent the ancient population groupings for which we could obtain dates of admixture (adjusted for sample date by assuming 28 years per generation): light green, 1400 BP; purple, 200 BP; red, 150 BP. Bars show one standard error in each direction. See **Table S4** for full results.

## Phylogeny of First Remote Oceanian ancestry

To test whether the First Remote Oceanian ancestry in ancient and present-day groups is more closely related to Lapita samples from Tonga or Vanuatu, we used a block jackknife to evaluate the difference between the statistics *f*_*4*_(*Test*, Han; Atayal, Tonga_2600BP) and *f*_*4*_(*Test*, Han; Atayal, Vanuatu_3000BP) for Oceanian populations as *Test* (STAR Methods). We found a trend toward greater allele-sharing with Tonga, with significant results in Polynesian and to a lesser degree Polynesian outlier populations (**Table S5**). These results show that the First Remote Oceanian ancestry in Polynesians today is derived from a source that was closer to the sampled Lapita-period population from Tonga than to the Vanuatu Lapita population. We do not observe significant differences for present-day populations from Vanuatu, but our statistical power is limited due to the small proportions of First Remote Oceanian ancestry.

## Phylogeny of Papuan ancestry

We built admixture graphs to explore in more detail the different streams of Papuan ancestry present in Oceania. We used as reference populations Australia, Kankanaey, Atayal, and Mixe, together with representatives of major poles of Papuan genetic variation inferred from the ADMIXTURE analysis: Vanuatu_Tanna, Mamusi (New Britain), Nasioi (Solomon Islands), New Guinea, and Tolai (New Britain/New Ireland). To avoid overfitting, we adopted a restricted framework in which the ancestry in each population was modeled as a combination of the same set of source lineages, with the exception of the unadmixed New Guinea population. We found that three Papuan source lineages were necessary in order to obtain a good fit for the model—one maximized in Mamusi, one maximized in Nasioi, and one closest to New Guinea—showing that the implied ancestry components from ADMIXTURE (**Figure S1**) are all well-supported in formal models based on allele-sharing statistics (**Figure S3**). The admixture graph analysis suggests that the blue (Bismarck Archipelago-majority) and red (Solomon Islands-majority) ADMIXTURE components represent admixed ancestry: both include First Remote Oceanian ancestry (~20% for red and ~5% for blue), and the two are additionally admixed with each other, as we could not fit a Solomon Islands population (e.g., Nasioi) and a Bismarck Archipelago population (e.g., Mamusi or Baining) simultaneously without admixture from one to the other. In our models, we included Solomon Islands-type ancestry in Mamusi (approximately one-third of its total Papuan ancestry), although we were unable to distinguish the direction(s) of gene flow. Vanuatu was confidently inferred to have ancestry from all three Papuan sources (Z > 8 for omitting any source).

We next asked if we could add Polynesians (Tongan) as a mixture of a component related to one of the other Oceanian populations along with additional First Remote Oceanian ancestry. Such a model was successful only in one configuration, with Tongan as a mixture of Tolai-related and First Remote Oceanian ancestry (all *f*-statistics fit to within 2.0 standard errors of their observed values except for one residual, *f*_*4*_(Kankanaey, Tongan; Australian, Vanuatu_Tanna), at Z = 2.7; **Figure 3** **and Figure S3**). Our choice to include Tolai in the model was guided by the ADMIXTURE analysis, in which the Papuan ancestry profile in Polynesians appears to match that in Tolai (and Tutuba, from near Espiritu Santo Island in Vanuatu) more closely than other populations. We note that the Tolai are known to be descended from relatively recent mixture between groups from New Ireland and New Britain (resulting from displacement caused by the eruption of the Rabaul caldera ~1400 BP [19]), so their ancestors cannot represent the true source population of the Papuan ancestry in Polynesians. However, the similarity of Tolai Papuan ancestry to Polynesians suggests that the Papuan component in Polynesians could similarly be from a mixture of multiple Near Oceanian sources. Given that Tolai are intermediate between populations from New Britain and New Ireland (the latter with high Solomon Islands-related ancestry), Polynesians could plausibly have acquired New Britain-related ancestry from Vanuatu or Santa Cruz, along with ancestry more closely related to that in New Ireland or the Solomon Islands via a distinct stream of migration.

As suggested by their similar mixtures of components in ADMIXTURE, the ancient Vanuatu individuals are broadly consistent with descent from the same common ancestral population as present-day groups from Vanuatu. In the admixture graphs, we could fit the ancient sample groups from 2400-200 BP as sister populations to Vanuatu_Tanna, albeit with different proportions of First Remote Oceanian ancestry. The one exception was the 150 BP grouping of individuals from Efate (with ~20% First Remote Oceanian ancestry), which showed significant un-modeled allele sharing with Tongan (max residual Z = 3.5, after accounting for excess First Remote Oceanian ancestry). Some present-day Vanuatu populations, such as Efate and Makura, show a similar pattern when added to the model, likely reflecting migration of Polynesians to Vanuatu in the last thousand years or less.

**Figure 3.**
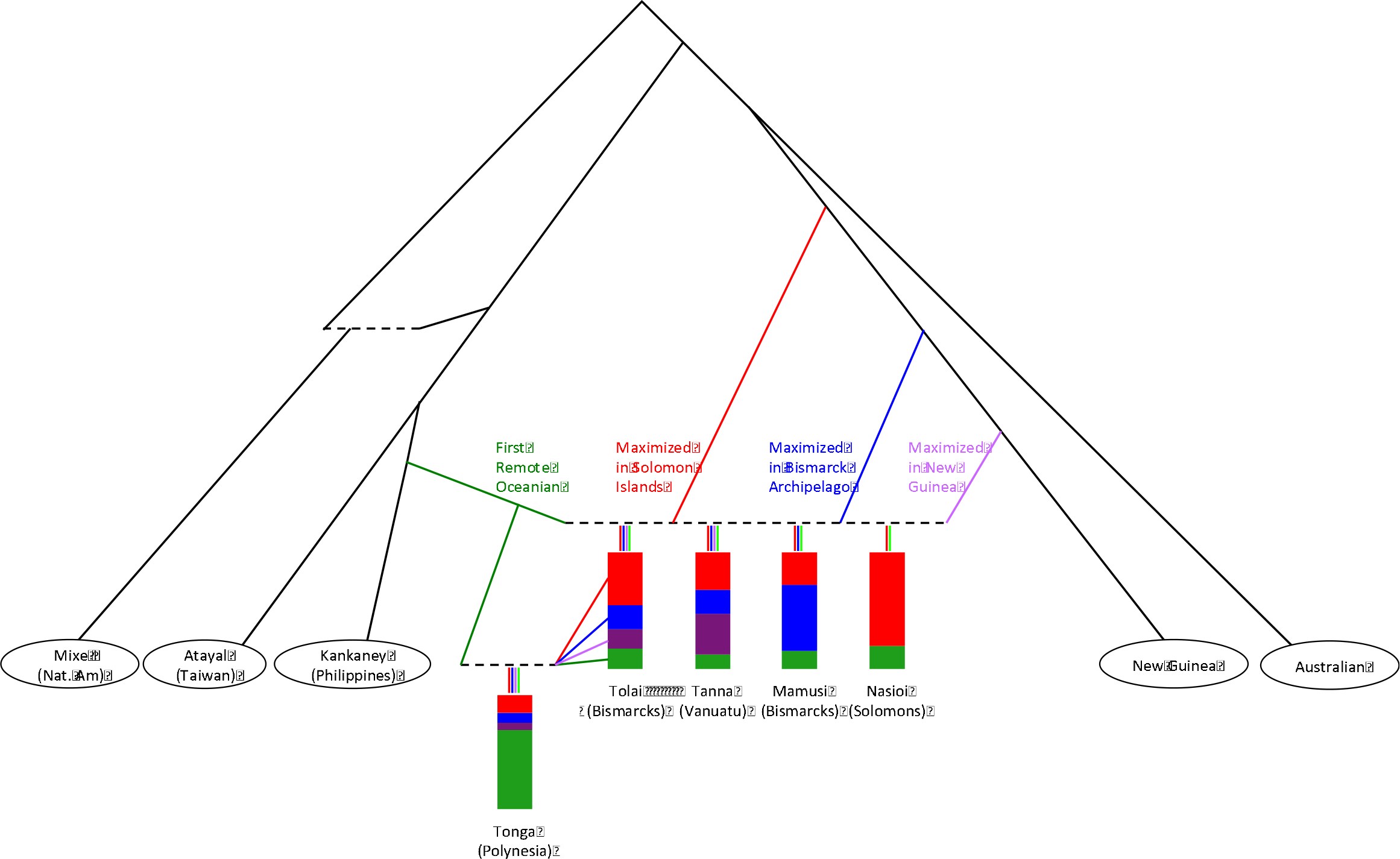
Working admixture graph model with diverse present-day Oceanian populations. Dotted lines denote admixture events. For five populations, the proportions of four fitted ancestry sources maximized in First Remote Oceanians (green), Solomon Islands (red), Bismarck Archipelago (blue) and New Guinea (purple) are shown. Papuan ancestry is inferred to be highly simlar in the Tolai and in Tonga, allowing Tonga to be fit as a mixture of a Tolai-related group and additional ancestry from First Remote Oceanians. We note that the colors are chosen to be correlated to the components inferred from ADMIXTURE (**Figure S1**), but the ADMIXTURE components represent combinations of the admixture graph sources given here, and hence the ratios differ between the two methods. Full model parameters can be found in **Figure S3**.

## Conclusion

By analyzing a time transect of Vanuatu from initial settlement through the present, combined with dense geographical sampling of surrounding present-day populations, we document a series of dramatic genetic shifts associated with consistently high human mobility through a total of at least four distinct streams of migration and admixture. First, the initial human migration to Vanuatu involved First Remote Oceanians associated with the Lapita culture. Second, by 2400 BP, these groups were almost completely displaced in Vanuatu by Papuan-ancestry populations originally from the Bismarck Archipelago, who remain the source for most of the ancestry of people in Vanuatu today. Third, in Polynesia, we find evidence for a different Papuan ancestry type that reflects a distinct migration. And fourth, finally, these streams of ancestry reconnected in parts of the Vanuatu archipelago, influenced by back-migration from Polynesia. These results highlight the importance of multiple episodes of migration and mixture in shaping the human diversity of Oceania.

## Acknowledgements

We are grateful to Fiona Petchey and Tomasz Goslar for sharing unpublished information on previously reported radiocarbon dates generated at the University of Waikato and the Poznan Accelerator Mass Spectrometry laboratories. The Teouma research was supported by the Australian Research Council (Discovery Grants DP0880789 and DP1101014 15, M.S. and S.B.), the National Geographic Society (M.S. and S.B.), the Australia-Pacific Science Foundation (M.S. and S.B.), the Royal Society of New Zealand Marsden Fund (UOO0917, H.B. and S.B.), and a University of Otago Research Grant (H.B.). We are grateful to the late Richard Shutler Jr. for access to his original field notes, and to David Burley for contributing further Shutler archives to the Vanuatu Cultural Centre which aided in interpretation. Ralph Regenvanu, former Director of the Vanuatu Cultural Centre, gave ethical guidance on the use of the present-day samples for this project. A.J.M. was supported by a Wellcome Trust Clinical Research Training Fellowship grant reference 106289/Z/14/Z. We thank Professors John Clegg, David Weatherall, Donald Bowden and their colleagues for their work establishing the Oceanic sample collection at the University of Oxford in the U.K, with support from the Wellcome Trust and Medical Research Council. F.V. was supported by CNRS-UMR 7041. P.S. was supported by the Swedish Research Council (VR grant 2014-453). Accelerator Mass Spectrometry radiocarbon dating work at Pennsylvania State University (D.J.K) was supported by the NSF Archaeometry program (BCS-1460369). D.R. was supported by NIH grant GM100233, by NSF HOMINID grant BCS-1032255, and by an Allen Discovery Center of the Paul Allen Foundation, and is a Howard Hughes Medical Institute investigator.

## Author Contributions

R.P. and D.R. supervised the study. M.S., F.V., S.B., R.S., H.B., I.P., G.W., and R.P. provided ancient samples and assembled archaeological and anthropological information. N.R., N.B., O.C., M.F., M.M., J.O., K.Si., and K.St. performed ancient DNA laboratory work. T.K.H. and D.J.K. carried out and analyzed radiocarbon dating data. K.A., A.H., K.M., S.J.O., T.P., K.R., T.N.W., and A.J.M. provided data from present-day populations. M.L., P.S., S.M., and D.R. analyzed genetic data. M.L., P.S., M.S., and D.R. wrote the manuscript.

## STAR Methods

### CONTACT FOR REAGENT AND RESOURCE SHARING

Further information and requests for resources and reagents should be direct to and will be fulfilled by the Lead Contact, David Reich

### EXPERIMENTAL MODEL AND SUBJECT DETAILS

**Archaeological Context on Ancient Individuals with New Genome-Wide Data.** We newly report data from 14 ancient skeletons. For 3 of these skeletons we are reporting new ancient DNA data increasing the quality of the dataset beyond what was reported on the samples in a previous study [1]. For the 11 remaining samples the data are entirely new:

#### Teouma, Efate Island (~3000 BP) – Lapita Culture (n=4 samples)

The Teouma Lapita culture cemetery and settlement site is discussed in detail in the Supplementary Information to Skoglund et al. 2016 and references [1]. The additional sample I5951 was displaced during quarrying activities before controlled archaeological excavations began at the site in 2004. Given its age it was highly likely to have been from a disturbed burial context of Lapita age and can be legitimately considered with the other Lapita-age skeletons from the site.

- I5951 (TeoQE), Vanuatu_3000BP

Newly reported sample
Genetic Sex: Male
Radiocarbon Date: 1258-1088 calBCE (2955±20, PSUAMS-2411)
- I1370_all (B17.P3), Vanuatu_3000BP

Previously reported in [1]; here we report higher coverage data
Genetic Sex: Female
Radiocarbon Date: 1160-830 calBCE (3083±26 BP, Wk-21026, corrected for Marine Reservoir Effect [1])
- I1369_all (B10B.P3), Vanuatu_3000BP

Previously reported in [1]; here we report higher coverage data
Genetic Sex: Female
Radiocarbon Date: 1050-800 calBCE (3045±30 BP, Poz-81126, corrected for Marine Reservoir Effect [1])
- I1368_all (TB30A.P3), Vanuatu_3000BP

Previously reported in [1]; here we report higher coverage data
Genetic Sex: Female
Radiocarbon Date: 1040-790 calBCE (2983±32 BP, Wk-22657, corrected for Marine Reservoir Effect [1])

#### Mele-Taplins, Efate Island (~2400 BP) (n=1 sample)

The Mele-Taplins site is described by Valentin and colleagues [20]. The skeleton comes from a subsurface grave in a rockshelter (Taplins 1) at the base of a cliff, excavated by Graeme Ward of The Australian National University in 1973-4 and curated at Otago University, Dunedin, New Zealand. Other burials from the Taplins 2 shelter were of broadly similar age.

- I4451_all (TAP1), Vanuatu_2400BP

Newly reported sample
Genetic Sex: Male
Radiocarbon Date: 516-369 calBCE (2348±32 BP, Wk-20390)

#### Burumba, Epi Island (~1400 BP) (n=2 samples)

The Burumba site is described by Valentin and colleagues [20] and excavated in 2006 by Frederique Valentin and Jacques Bole. The graves of nine adults were excavated from an open site at Kalala Plantation 200m from the current beach, dug into sterile sand. Burial 5 was an assemblage of cranial remains of five individuals placed on a pile of coral slabs and blocks.

- I3921_all (BURU5D), Vanuatu_1400BP

Newly reported sample
Genetic Sex: Male
Radiocarbon Date: 429-663 calCE [429-595 calCE (1530±20 BP, PSUAMS-1841), 619-663 calCE (1395±15, PSUAMS-2428)]
- I4096_all (BURU5B), Vanuatu_1400BP

Newly reported sample
Genetic Sex: Male
Radiocarbon Date: 545-650 calCE [551-650 calCE (1464±30 BP, Wk-25769), 545-610 calCE (1490±15 BP, PSUAMS-2460)]

#### Mangaliliu, Efate Island (~600 BP) (n=1 sample)

The burial was excavated from a test pit in Mangaliliu village by Richard Shing in 2002 and published in detail by Valentin and colleagues [21]. The originally reported age of the burial was reassessed after direct dating of the skeleton [20].

- I5259 (burial 1, Mang1), Vanuatu_600BP, Mangaliliu (Efate Island)

Newly reported sample
Genetic Sex: Female
Radiocarbon Date: 1307-1430 calCE (559±30 BP, Wk-20030)

#### Pangpang, Efate Island (~350 BP) (n=1 sample)

This burial, in a flexed position, was excavated by Richard Shing and Iarawai Philip during archaeological impact assessment related to the Efate Ring Road construction, between the villages of Pangpang and Forari. The body was adorned with ornaments composed of numerous tiny Conus shell and shark vertebrae beads and a large pearl shell pendant. This range of ornaments has been recorded in burial contexts of the last 400 years, prior to and during the initial phases of European contact (unpublished field notes, Vanuatu National Museum).

- I4450 (SEPU1, Sepulture 1), Vanuatu_350BP

Newly reported sample
Genetic Sex: Female
Radiocarbon Date: 1520-1645 calCE (305±15 BP, UCIAMS-188793)

#### Wam Bay, Epi Island (~200 BP) (n=2 samples)

The site appears to have been a largely Mission period, late 19^th^ to early 20^th^ century, cemetery of which three burials were exposed and was in proximity to a combustion feature associated with the making of lime-plaster for construction, a European introduced practice. The date of these burials may need to be further calibrated in the light of dietary analysis and could be younger than indicated by current calibration of the bone dates. The site was excavated by Frederique Valentin and Matthew Spriggs in 2006 (unpublished field notes, Vanuatu National Museum).

- I4105_all (WAMB1), Vanuatu_200BP

Newly reported sample
Genetic Sex: Male
Radiocarbon Date: 1529-1798 calCE (255±20 BP, PSUAMS-1922)
- I4106_all (WAMB2), Vanuatu_200BP

Newly reported sample
Genetic Sex: Male
Radiocarbon Date: 1645-1950 calCE (225±20 BP, PSUAMS-1923)

#### Ifira, Efate Island (Historical Period) (n=1 sample)

This tightly flexed burial from a feature containing skeletal remains of two individuals was excavated by Mary Elizabeth and Richard Shutler, Jr, in June 1964 on the small island of Ifira in Vila Harbor, Port Vila, during a test pit survey of the island. It is briefly mentioned in Shutler and Shutler [22]. Unpublished field notes relating to the excavation are held in the files of the Vanuatu National Museum. Ifira is notable as one of the Vanuatu Polynesian Outlier islands and this burial would date to the period of Polynesian cultural influence.

- I4425 (EF3_2_E, Pit 2; Loc E), Vanuatu_150BP

Newly reported sample
Genetic Sex: Female
Radiocarbon Date: 1652-1950 calCE (200±20 BP, UCIAMS-188795)

#### Pango Village, Efate Island (Historical Period) (n=1 sample)

This is one of two individuals excavated by Mary Elizabeth and Richard Shutler, Jr, in June 1964 on the Pango Peninsula opposite the small island of Ifira in Vila Harbour, Port Vila. Unpublished field notes relating to the excavation are held in the files of the Vanuatu National Museum, but little detail is available.

- I4424 (EF_Pango1), Vanuatu_150BP

Newly reported sample
Genetic Sex: Male
Radiocarbon Date: 1661-1950 calCE (190±15 BP, UCIAMS-188794)

#### Banana Bay, Efate Island (Historical Period) (n=1 sample)

The burial was excavated by Richard Shing and Iarawai Philip during archaeological impact assessment related to the Efate Ring Road construction in the Banana Bay area, southeast Efate. The body, lying on the back, was adorned with ornaments including numerous tiny Conus shell beads and a few European glass beads (unpublished field notes, Vanuatu National Museum).

- I4419 (BB1, Burial 1), Vanuatu_150BP, Banana Bay (Efate Island)

Newly reported sample
Genetic Sex: Male
Radiocarbon Date: 1678-1940 calCE (135±15 BP, UCIAMS-188792)

**Data Collection Strategy for Newly Reported Data from Present-Day Vanuatu.** We genotyped 185 present-day individuals from 32 populations from Vanuatu spanning 18 islands. All individuals gave informed verbal consent for studies of population history and human health, especially anemia, consistent with the standards prevailing at the time the data were collected. Samples of whole blood were collected as part of a range of research projects undertaken from the late 1970s in collaborations between multiple sites and institutions in Vanuatu and the University of Oxford investigating population differences at the genetic level. In accordance with participant consent, DNA was extracted, anonymized, and stored in batches analyzable only by geographic location of participant origin. Use of the samples for genome-wide analyses including studies of population history was reviewed by the Oxford Tropical Research Ethics Community at the University of Oxford and formally approved in a letter dated July 2 2014 (OXTREC Reference: 537-14). The use of the samples for genetic analysis was also approved by the Vanuatu Cultural Centre in a formal letter dated May 30, 2017.

## METHOD DETAILS

**Ancient DNA laboratory work.** In a dedicated clean room at University College Dublin, we used a dental sandblaster to separate cochlear sections from petrous bones. We milled these samples into fine powder, and shipped them to Harvard Medical School.

At Harvard Medical School, we extracted DNA following a previously published protocol [4], with two modifications. First, we replaced the combination of a funnel and a MinElute column with Roche columns [5]. Second, we eluted two times in 45μl, obtaining 90μl of extract for each sample.

We prepared libraries from the extracts using a double-stranded protocol, affixing 7-base-pair sequences to either end to allow multiplexing of the libraries and to prevent contamination from affecting the samples after barcodes were added. We prepared some of the libraries in the presence of the enzyme UDG to remove characteristic damage associated with ancient DNA (**Table S2**) [6].

We enriched the libraries in solution for sequences overlapping the mitochondrial genome [8] as well as for 3000 nuclear positions, and sequenced on an Illumina NextSeq500 instrument for 2×76cycles + 2×7 cycles after adding a pair of unique 7-base-pair indices. For libraries that were promising after screening, we next enriched for sequences overlapping approximately 1.24 million SNPs [9, 23–25]. We added unique 7-base-pair index combinations to each enriched library, and sequenced on a multiplexed pool of samples on a lane of an Illumina NextSeq500 instrument for 2×76cycles + 2×7cycles. We iteratively sequenced more sequences from each sample until the number of new SNPs covered per additional sequences generated was less than about 1 in 100.

For samples for which we wished to obtain more coverage, we prepared additional libraries from existing extract or new extract, up to 8 libraries for some samples. We pooled data from all libraries for further analysis.

**Bioinformatic processing.** We demultiplexed reads into libraries based on their two indices and two barcodes, allowing no more than one mismatch to the total of four expected 7 base pair sequences. We merged sequences requiring at least 15 base pairs of overlap using *SeqPrep* (github.com/jstjohn/SeqPrep).

We aligned merged sequences to the mitochondrial RSRS genome [26] (for mitochondrial DNA analyses) and to the hg19 reference (for whole genome analyses). For alignment we used the single-ended aligner “samse” from BWA with default parameters (version 0.6.1) [27]. For samples which are non-UDG treated (and therefore may have higher mismatch rates compared to the reference genome), we used more relaxed alignment parameters, “-n 0.01 -o 2 -l 16500”. This setting disables seeding, allowing for less conservative alignments, helping to align damaged reads.

**Haplogroup calling strategy on mitochondrial DNA data.** We determined haplogroups using Haplogrep2, which provides a reliability score for assigned haplogroups [28]. We ran Haplogrep2 in three configurations and picked the best rank score to represent the haplogroup for that individual. (a) We restricted sequences to those with characteristic patterns of ancient DNA damage in their terminal nucleotides, which removes contamination. To do this, we used the PMDtools software [29] requiring a minimum score of pmdscore=3. We trimmed the sequences obtained in this way by 5 base pairs on either side to remove nucleotides likely to be deaminated prior to running Haplogrep2. (b) As a second approach, we trimmed sequences by 5 base pairs on either side to eliminate characteristic ancient DNA damage and fed these sequences to Haplogrep2 without damage restriction. (c) Finally, we applied no trimming and made a haplogroup call. We manually made two exceptions to the rule of always picking the best ranking call. For S4106.E1.L1, (a) and (c) gave similar ranking scores and we selected B4a1a1a11 from method (a) based on consistency with calls from two other libraries from the same sample. For S4096.E1.L2, we selected B4a1a1k manually from method (a) despite a marginally lower rank score than method (c).

## QUANTIFICATION AND STATISTICAL ANALYSIS

### Population genetic analyses

All analyses were based on the set of 593,124 autosomal Human Origins SNPs, except for ADMIXTURE, which was performed with all 597,573 Human Origins SNPs. Principal component analysis was carried out using the “lsqproject” and “autoshrink” options in smartpca [30, 31]. ADMIXTURE [10] clustering analysis was performed using default parameters, with the cluster components (K) ranging from K=2 to K=8. *f*-statistics were computed in ADMIXTOOLS [32], using the qp4diff program for differences between Lapita *f*_*4*_-statistics (“allsnps” mode), with standard errors obtained by block jackknife.

### Admixture graph fitting

We constructed admixture graphs using the qpGraph utility in ADMIXTOOLS [32]. Mixe’s position as an outgroup relative to the other populations (in an unrooted sense) means that its eastern and western Eurasian ancestry components can be collapsed into a single lineage with no change in the model. Similarly, we can omit explicit inclusion of Denisovan admixture because of the symmetry of such ancestry in the right-hand clade of the model (as displayed in **Figure S3**).

## DATASET AND SOFTWARE AVAILABILITY

Raw sequences from the 14 individuals are available from the European Nucleotide Archive at accession number PRJEB24938. Genotypes are available at https://reich.hms.harvard.edu/datasets. To access data for the newly genotyped present-day individuals from Vanuatu, researchers should send a signed letter to D.R. containing the following text: “(a) I will not distribute the data outside my collaboration; (b) I will not post the data publicly; (c) I will make no attempt to connect the genetic data to personal identifiers for the samples; (d) I will use the data only for studies of population history; (e) I will not use the data for any selection studies; (f) I will not use the data for medical or disease-related analyses; (g) I will not use the data for commercial purposes.”

## KEY RESOURCES TABLE

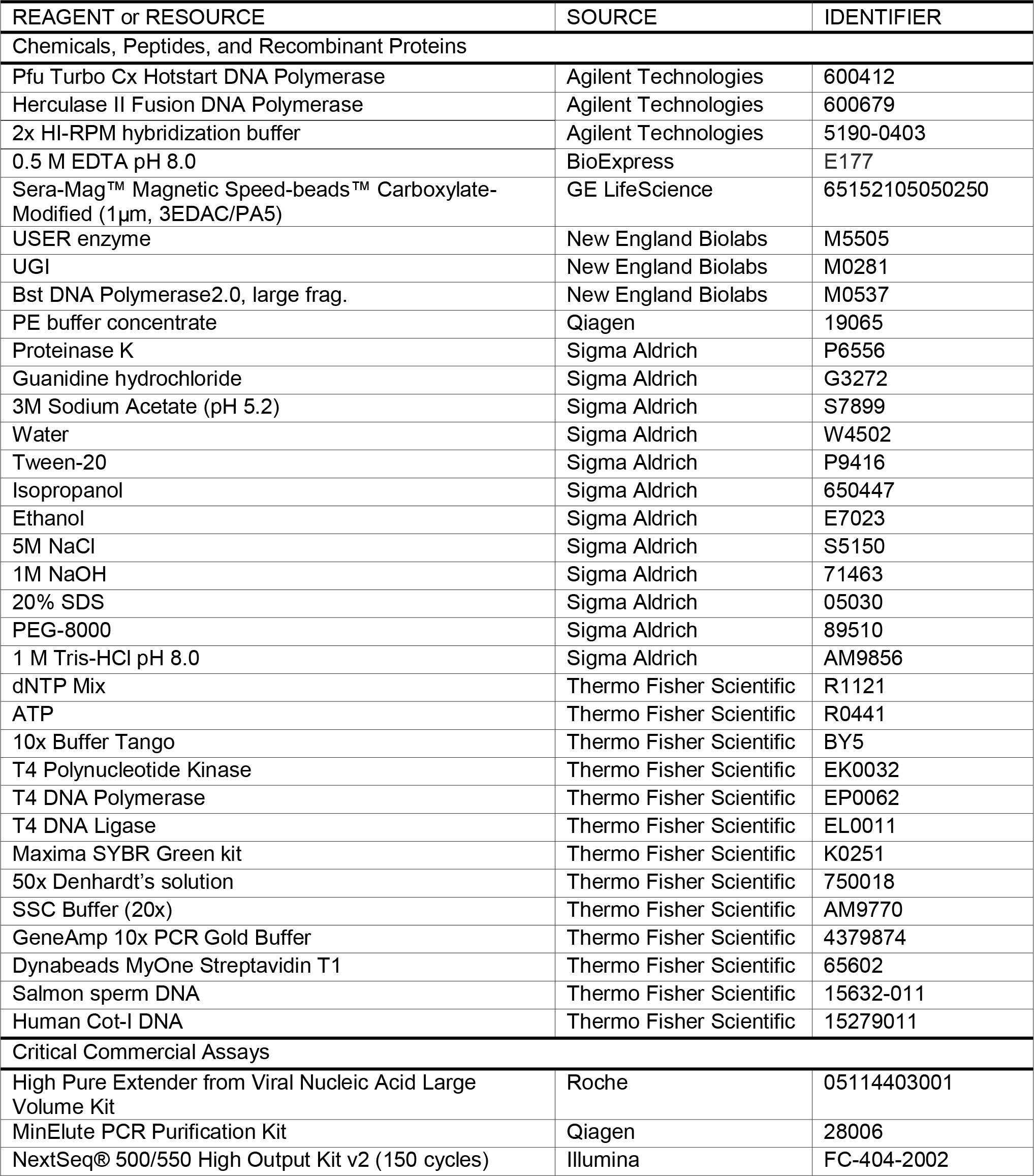

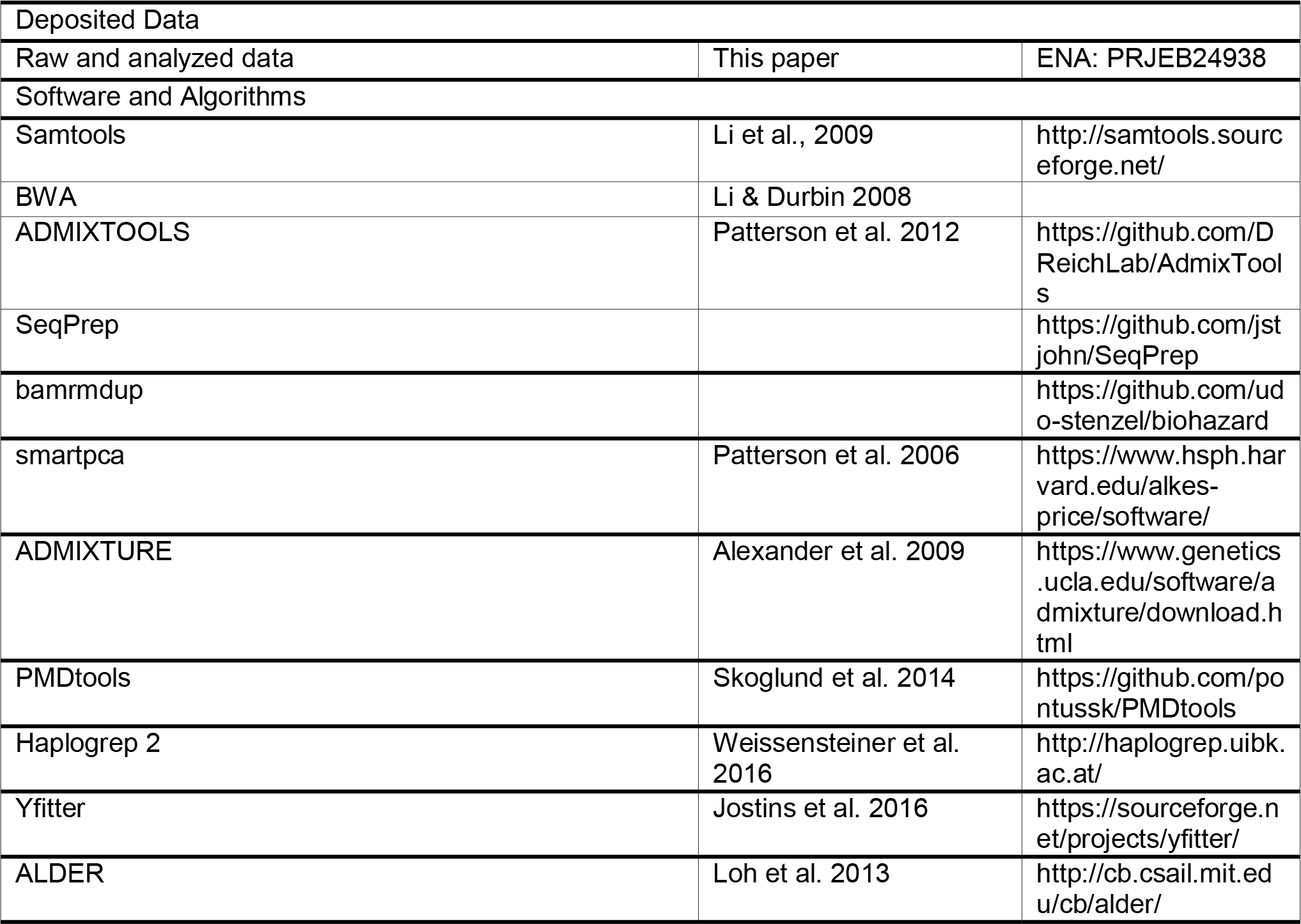

---

**Figure S2.**
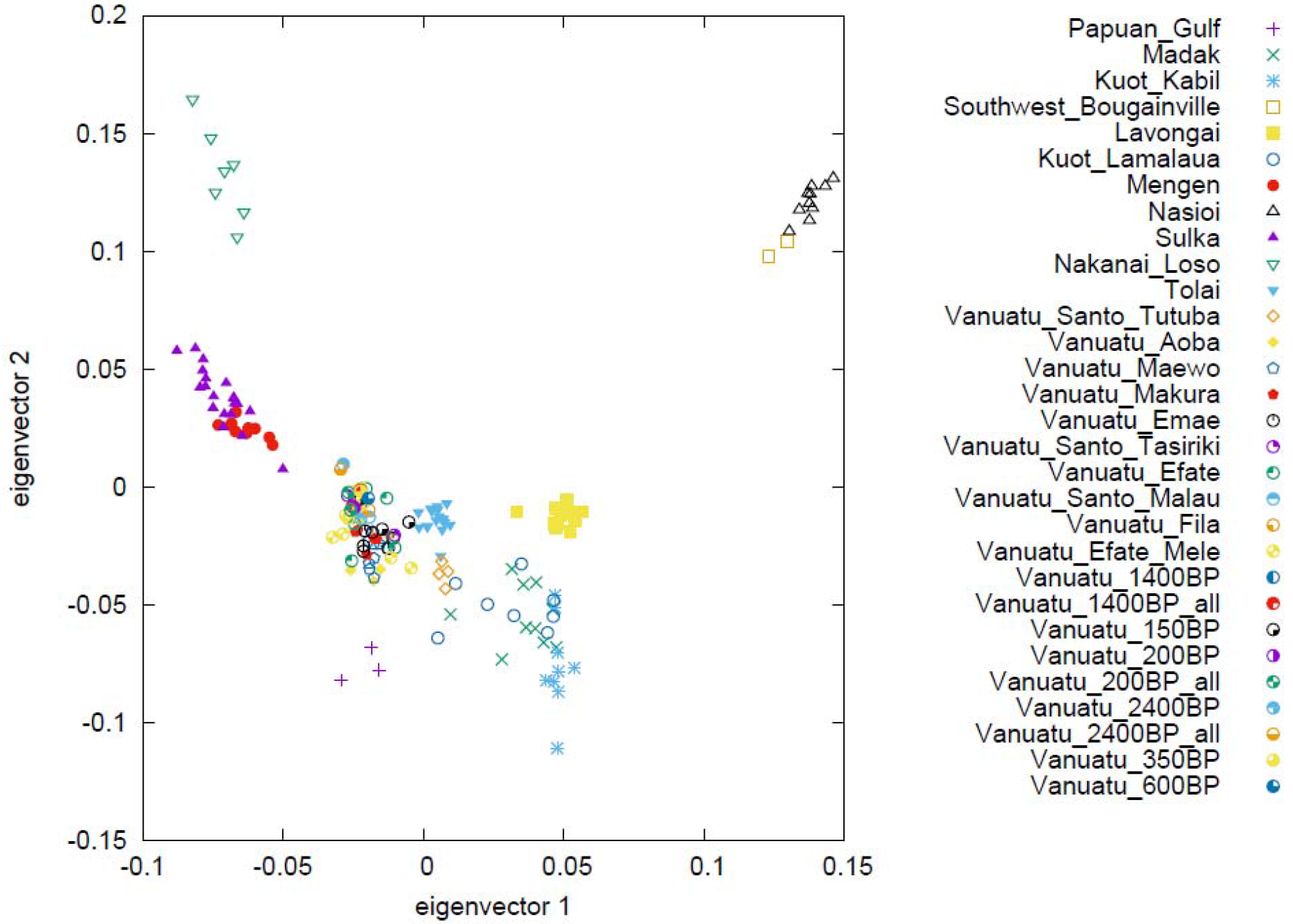
Principal component analysis of Oceanian populations. We computed axes using present-day populations with 17-25% First Remote Oceanian ancestry and projected ancient samples. For samples with a combination of partial-UDG-treated and non-UDG libraries, the combined data (“_all”) are very similar to the UDG-only data, which enhances our confidence in the results.

**Figure S3.**
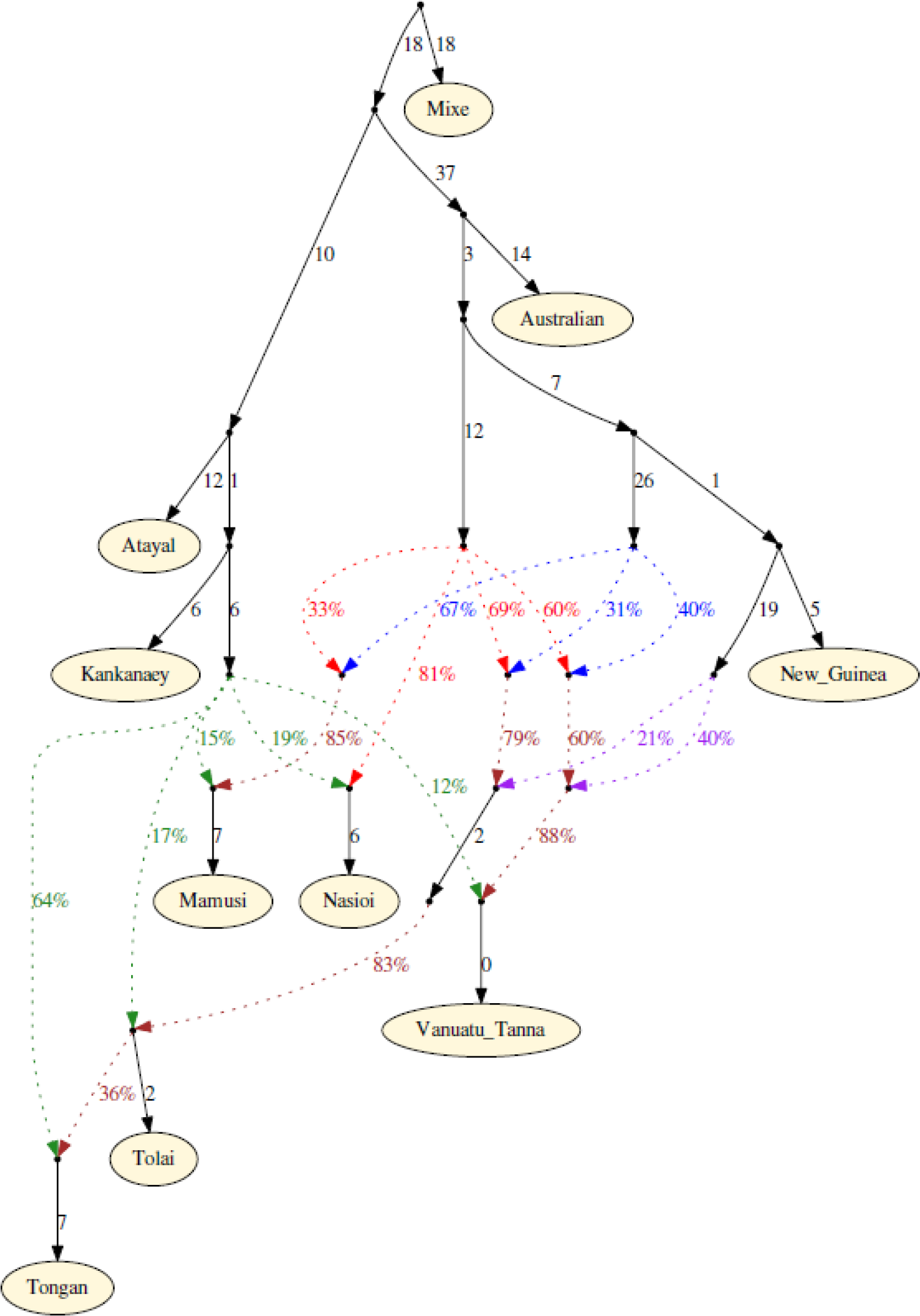
Admixture graph model with inferred parameters. The model shown is the same as in **Figure 3** but with an alternative visualization. Branch lengths are given in units of *f*_*2*_ genetic drift distance times 1000, and admixture proportions are indicated along corresponding dotted lines. Red, Solomon Islands majority source; blue, Bismarck Archipelago majority source; purple, New Guinea-related source; green, First Remote Oceanian; brown, mixed ancestry. The order of admixture events specified is arbitrary.

## References

1. Skoglund, P., Posth C., Sirak K., Spriggs M., Valentin F., Bedford S., Clark, G.R., Reepmeyer C., Petchey F., Fernandes D., et al. (2016). Genomic insights into the peopling of the Southwest Pacific. Nature 538, 510–513.

2. Valentin F., Detroit F., Spriggs, M.J., and Bedford S. (2016). Early Lapita skeletons from Vanuatu show Polynesian craniofacial shape: Implications for Remote Oceanic settlement and Lapita origins. Proceedings of the National Academy of Sciences of the United States of America 113 292–297.

3. Pinhasi R., Fernandes D., Sirak K., Novak M., Connell S., Alpaslan-Roodenberg, S., Gerritsen F., Moiseyev V., Gromov A., Raczky P., et al. (2015). Optimal Ancient DNA Yields from the Inner Ear Part of the Human Petrous Bone. PloS one 10 e0129102.

4. Dabney J., Knapp M., Glocke I., Gansauge, M.T., Weihmann A., Nickel B., Valdiosera C., Garcia N., Paabo S., Arsuaga, J.L., et al. (2013). Complete mitochondrial genome sequence of a Middle Pleistocene cave bear reconstructed from ultrashort DNA fragments. Proceedings of the National Academy of Sciences of the United States of America 110 15758–15763.

5. Korlevic P., Gerber T., Gansauge, M.T., Hajdinjak M., Nagel, S., Aximu-Petri, A., and Meyer M. (2015). Reducing microbial and human contamination in DNA extractions from ancient bones and teeth. BioTechniques 59 87–93.

6. Rohland N., Harney E., Mallick S., Nordenfelt S., and Reich D. (2015). Partial uracil-DNA-glycosylase treatment for screening of ancient DNA. Philosophical transactions of the Royal Society of London. Series B, Biological sciences 370 20130624.

7. Briggs, A.W., Stenzel U., Meyer M., Krause J., Kircher M., and Paabo S. (2010). Removal of deaminated cytosines and detection of in vivo methylation in ancient DNA. Nucleic acids research 38 e87.

8. Maricic T., Whitten M., and Paabo S. (2010). Multiplexed DNA sequence capture of mitochondrial genomes using PCR products. PloS one 5 e14004.

9. Fu Q., Meyer M., Gao X., Stenzel U., Burbano, H.A., Kelso J., and Paabo S. (2013). DNA analysis of an early modern human from Tianyuan Cave, China. Proceedings of the National Academy of Sciences of the United States of America 110 2223–2227.

10. Alexander, D.H., Novembre J., and Lange K. (2009). Fast model-based estimation of ancestry in unrelated individuals. Genome research 19 1655–1664.

11. Pugach I., Duggan, A.T., Merriwether, D.A., Friedlaender, F.R., Friedlaender, J.S., and Stoneking M. (2018). The gateway from Near into Remote Oceania: new insights from genome-wide data. Molecular biology and evolution.

12. Reich D., Thangaraj K., Patterson N., Price, A.L., and Singh L. (2009). Reconstructing Indian population history. Nature 461 489–494.

13. Moorjani P., Patterson N., Hirschhorn, J.N., Keinan A., Hao L., Atzmon G., Burns E., Ostrer H., Price, A.L., and Reich D. (2011). The history of African gene flow into Southern Europeans, Levantines, and Jews. PLoS genetics 7 e1001373.

14. Loh, P.-R., Lipson M., Patterson N., Moorjani P., Pickrell, J.K., Reich D., and Berger B. (2012). Inference of admixture parameters in human populations using weighted linkage disequilibrium.

15. Moorjani P., Sankararaman S., Fu Q., Przeworski M., Patterson N., and Reich D. (2016). A genetic method for dating ancient genomes provides a direct estimate of human generation interval in the last 45,000 years. Proceedings of the National Academy of Sciences of the United States of America 113 5652–5657.

16. Garanger J. (1972). Archéologie des Nouvelles-Hébrides, (Paris: ORSTOM).

17. Xu S., Pugach I., Stoneking M., Kayser M., Jin L., and Consortium, H.P.-A.S. (2012). Genetic dating indicates that the Asian-Papuan admixture through Eastern Indonesia corresponds to the Austronesian expansion. Proceedings of the National Academy of Sciences of the United States of America 109 4574–4579.

18. Lipson M., Loh, P.R., Patterson N., Moorjani P., Ko, Y.C., Stoneking M., Berger B., and Reich D. (2014). Reconstructing Austronesian population history in Island Southeast Asia. Nature communications 5 4689.

19. Spriggs M. (1997). The Island Melanesians, (London: Routledge).

20. Valentin F., Spriggs M., Bedford S., and Buckley H. (2011). Vanuatu mortuary practices over three millennia: Lapita to the early contact period. Journal of Pacific Archaeology 2 49–65.

21. Valentin F., Shing R., and Spriggs M. (2005). Des restes humains datés du début de la période de Mangaasi (2400-1800 BP) découverts à Mangaliliu (Efate, Vanuatu). Comptes Rendus Palé;vol 4 420–427.

22. Shutler, M.E., and Shutler R. (1966). A preliminary report of archaeological explorations in the Southern New Hebrides. Asian Perspectives 9 157–166.

23. Fu Q., Hajdinjak M., Moldovan, O.T., Constantin S., Mallick S., Skoglund P., Patterson N., Rohland N., Lazaridis I., Nickel B., et al. (2015). An early modern human from Romania with a recent Neanderthal ancestor. Nature 524 216–219.

24. Haak W., Lazaridis I., Patterson N., Rohland N., Mallick S., Llamas B., Brandt G., Nordenfelt S., Harney E., Stewardson K., et al. (2015). Massive migration from the steppe was a source for Indo-European languages in Europe. Nature 522 207–211.

25. Mathieson I., Lazaridis I., Rohland N., Mallick S., Patterson N., Roodenberg, S.A., Harney E., Stewardson K., Fernandes D., Novak M., et al. (2015). Genome-wide patterns of selection in 230 ancient Eurasians. Nature 528 499–503.

26. Behar, D.M., van Oven, M., Rosset S., Metspalu M., Loogvali, E.L., Silva, N.M., Kivisild T., Torroni A., and Villems R. (2012). A “Copernican” reassessment of the human mitochondrial DNA tree from its root. American journal of human genetics 90 675–684.

27. Li H., and Durbin R. (2010). Fast and accurate long-read alignment with Burrows-Wheeler transform. Bioinformatics 26 589–595.

28. Weissensteiner H., Pacher D., Kloss-Brandstatter, A., Forer L., Specht G., Bandelt, H.J., Kronenberg F., Salas A., and Schonherr S. (2016). HaploGrep 2: mitochondrial haplogroup classification in the era of high-throughput sequencing. Nucleic acids research 44 W58–63.

29. Skoglund P., Northoff, B.H., Shunkov, M.V., Derevianko, A.P., Paabo S., Krause J., and Jakobsson M. (2014). Separating endogenous ancient DNA from modern day contamination in a Siberian Neandertal. Proceedings of the National Academy of Sciences of the United States of America 111 2229–2234.

30. Patterson N., Price, A.L., and Reich D. (2006). Population structure and eigenanalysis. PLoS genetics 2 e190.

31. Galinsky, K.J., Bhatia G., Loh, P.R., Georgiev S., Mukherjee S., Patterson, N.J., and Price, A.L. (2016). Fast Principal-Component Analysis Reveals Convergent Evolution of ADH1B in Europe and East Asia. American journal of human genetics 98 456–472.

32. Patterson N., Moorjani P., Luo Y., Mallick S., Rohland N., Zhan Y., Genschoreck T., Webster T., and Reich D. (2012). Ancient admixture in human history. Genetics 192 1065–1093.

